# A CD40-Targeted IL-21 Fusokine Enables Rapid Generation of Human IL-10⁺Granzyme B⁺ Regulatory B Cells

**DOI:** 10.64898/2026.01.12.699061

**Authors:** Andrea Pennati, Shala Yuan, Rahul Das, Catigan Hedican, Amar Yeware, Jacques Galipeau

## Abstract

Regulatory B cells (Bregs) have emerged as important modulators of immune homeostasis, complementing the classical roles of B cells in antibody production, antigen presentation, and provision of costimulatory signals and cytokines. Beyond promoting immunity, B cells can actively suppress inflammatory responses through specialized regulatory programs that limit effector T cell expansion and restrain pathogenic myeloid activation. These suppressive functions are mediated by mechanisms including secretion of IL-10, IL-35, and TGF-β, expression of inhibitory ligands, and release of granzyme B. However, Bregs are rare and heterogeneous in humans, typically comprising less than 1–2% of circulating B cells, and defects in their frequency or function are associated with autoimmune and inflammatory diseases.

The scarcity of endogenous Bregs has driven interest in their ex vivo generation as a cellular immunotherapy. Although several strategies efficiently induce regulatory B cells in murine systems, translation to human B cells has been limited by inefficiency, lack of scalability, or reliance on poorly defined culture conditions. Consequently, a robust and clinically compatible approach for generating human Bregs has remained elusive.

With human peripheral blood B cells as starting materiel, we show that a CD40 targeted IL-21 gain of function fusion protein in combination with TLR9 activation induces a stable regulatory phenotype characterized by IL-10 and granzyme B expression in more than 95% of input B-cells. These induced Bregs display a coordinated transcriptional program distinct from conventional activation, significantly suppress activated human T cell proliferation and modulate inflammatory myeloid responses in vitro. In mice with human T-cell driven xenoGVHD, transfusion of T-cell donor matched Bregs significantly improves survival. These data support the feasibility of Breg manufacturing at scale for use a cellular pharmaceutical.

## Introduction

B cells contribute to protective immunity through antibody production, antigen presentation, and provision of costimulatory signals and cytokines that shape T-cell and innate responses [1–3]. Nevertheless, the notion that B cells can also suppress the immune response it is not new [4]. This suppressive activity is attributed, at least in part, to heterogeneous populations collectively referred to as regulatory B cells (Bregs), which can limit effector T-cell expansion and inflammatory myeloid activation through soluble mediators and contact-dependent pathways [5–10]. Bregs, exert a negative role via the production of regulatory cytokines like IL-10, TGF-β, and IL-35 and the ability to express inhibitory molecules that suppress activated T cells and autoreactive B cells in a cell-cell contacted manner [11–13].

Naturally occurring Bregs represent less than 2% of all circulating blood lymphoid cells [14, 15] and abnormalities in the number and/or function of Bregs are found in several autoimmune diseases, including systemic lupus erythematosus (SLE), rheumatoid arthritis (RA), multiple sclerosis (MS), and psoriasis [16–20]. Given the rarity of Bregs in circulation, *ex vivo* expansion of B cells with regulatory attributes, is of great interest in the field of cell-therapy since pharmaceutical repletion of functional Bregs could be beneficial for the treatment of immune-mediated pathologies. A broad range of murine models have validated the therapeutic properties of Bregs [21–24]. *In vitro*, different stimuli can induce the conversion of B cells into IL-10 producing B cells with a regulatory phenotype. This includes microbial products as TLR4 and TLR9 ligands (LPS and CpG, respectively), or inflammatory cytokines such as IL-6, IL-1β and IFN-α as well CD40 ligation methods allow the differentiation of IL-10 producing B cells with a regulatory phenotype [15, 25–27]. Nevertheless, there is virtually no practical method to generate a clinically relevant amount of Bregs to meet cellular therapy demand, which informs the need for novel methods to produce sufficient Bregs compliant with Good Manufacturing Practice guidelines in anticipation of the Food and Drug Administration (FDA)-sanctioned first-in-human clinical trial.

Here, we demonstrate that uncommitted human peripheral blood B cells can be efficiently programmed in vitro into a regulatory state using a defined, cocultured-free approach. Stimulation with a CD40-targeted IL-21 gain-of-function fusion protein (CD40scFv–IL-21) in combination with a TLR9 agonist (CpG) induces a phenotypically distinct CD19⁺CD25⁺CD71⁺CD73⁻ B-cell population that expresses IL-10 and granzyme B. Transcriptomic profiling across independent healthy donors reveals a coordinated regulatory gene program that is distinct from CpG-driven activation alone. Functionally, these induced regulatory B cells suppress activated human T-cell proliferation and modulate inflammatory responses in monocyte-derived macrophages in vitro, and attenuate T cell–mediated disease progression in in a murine model of xenoGvHD. Together, these findings establish a robust and scalable strategy for generating human regulatory B cells from peripheral blood, supporting their further evaluation as a candidate cellular immunotherapy for immune-mediated disorders.

## Materials and Methods

### Ethics statement

All animal procedures followed the guidelines and recommendations of the Research Animal Resources Center and the Animal Care and Use Committee at the University of Wisconsin–Madison (IACUC M006547). NSG mice (NOD.Cg-Rag^1tm1Mom^*Il2rg*^tm1Wjl^/SzJ, strain #:007799) were obtained from The Jackson Laboratory. Anesthesia was administered using isoflurane, and euthanasia was performed via by CO_2_ asphyxiation.

All human research studies were performed in accordance with the Declaration of Helsinki. Prior to starting the study, ethical approval was obtained from the Institutional Review Board (IRB) at the University of Wisconsin – Madison (2019-0865-CP002).

### Mice and Genotyping

Mice were maintained under SPF conditions with ad libitum food and water. Male and female mice were used at 6–10 weeks of age. NSG genotype was verified using vendor documentation.

### ScFvCD40-IL-21 expression

Human embryonic kidney 293T cells stably expressing the CD40scFv–IL-21 fusion protein (ATCC CRL-3216) were cultured in FreeStyle™ 293 Expression Medium (Thermo Fisher Scientific, Cat. #12338-018) under suspension conditions. Briefly, frozen cell stocks were rapidly thawed and transferred into a 250 mL vented shake flask containing 30 mL of prewarmed medium. Cultures were maintained at 37 °C in a humidified atmosphere of 5% CO₂ on an orbital shaker at 160 rpm, with flask caps loosened to permit gas exchange.

Cells were expanded to a viable density of approximately 1 × 10⁶ cells/mL with viability ≥90% and routinely passaged at a 1:2 ratio every three days. For protein production, cells were cultured in 200–250 mL of medium in 500 mL shake flasks at 180 rpm. Culture supernatants containing secreted CD40scFv–IL-21 were harvested every two days and replaced with fresh medium.

Collected supernatants were clarified by centrifugation and concentrated using Amicon® Ultra centrifugal filter units (Millipore). Expression levels of the CD40scFv–IL-21 fusion protein were quantified by human IL-21 ELISA (BioLegend), according to the manufacturer’s instructions.

### PBMCs isolation and B/T/monocytes cells purification

PBMCs were isolated from heparinized blood of healthy human donors (n = 10) by Ficoll density gradient centrifugation. Briefly, anticoagulant-treated blood, diluted two-fold on DPBS, was carefully layered on top of one-volume of Ficoll-Paque PLUS density gradient media (GE Healthcare) in a tube. The tube was then centrifuged 400 xg for 30 min at 25 °C. This first centrifugation step generated three layers: the upper layer was plasma, the intermediate layer containing mononuclear cells and the bottom layer, which consisted mostly of erythrocytes and polymorphonuclear granulocytes. The plasma/Ficoll medium interface contains both PBMCs and platelets. To isolate PBMCs, this interface was carefully removed, washed with a salt-buffered solution, and then centrifuged. The supernatant, containing platelets, Ficoll and plasma, was removed, leaving a pellet of purified PBMCs.

Human B and T cells were purified from peripheral blood mononuclear cells using the EasySep™ Human B Cell Enrichment Kit or EasySep™ Human T Cell Enrichment Kit (BioLegend), following the manufacturer’s protocols. and monocyte by positive selection kit (StemSep™ Human CD14 Positive Selection Kit, STEMCELL Technologies), according to the manufacture instructions. Isolated cells were immediately cultured or cryopreserved or used for flow cytometry analysis. The purity of isolated B/T cells was routinely >95% as establish by flow cytometry analysis.

### Cell stimulation

Human B cells were cultured in complete R10 medium (RPMI-1640 with 10% fetal bovine serum-FBS (Sigma), 1 mM Sodium Pyruvate (Lonza), 2 mM L-Glutamine (Lonza), 10 mM HEPES (Lonza), 1% Essential Amino Acid (Gibco), 0.1 mM Beta-mercaptoethanol (Gipco) and the addition of 100 U/mL penicillin/streptomycin) and treated with 100 ng/mL CD40scFv–IL-21 and 0.1 μM CpG (ODN2006, Invivogen), or CpG only, in a 5% CO_2_ incubator for 3 to 5 days. Cells were then collected and spun down at 1,600 rpm for 5 min. Supernatant was collected for ELISA and cells were stained for phenotypic flow cytometry analysis. Alternatively, for T cells suppression assay and MDM co culture, B cells were stimulated for 24 h, then collected and washed twice with PBS and culture with T cells or MDM.

### Western blot analysis of CD40 and IL-21 signaling in human B cells

Human B cells were isolated from peripheral blood mononuclear cells (PBMCs) obtained from healthy donors using a BioLegend human B cell isolation kit, according to the manufacturer’s instructions. Cell purity was routinely ≥95% CD19⁺ as assessed by flow cytometry. Purified B cells were cultured in complete R10 medium and stimulated with purified CD40scFv–IL-21 (100 ng/mL), recombinant CD40 ligand (where indicated), or control conditions for the indicated time points.

Following stimulation, cells were harvested and lysed in ice-cold RIPA buffer supplemented with protease and phosphatase inhibitors for 20 min on ice. Lysates were clarified by centrifugation at 14,000 ×g for 15 min at 4 °C, and protein concentrations were determined using a BCA assay. Equal amounts of protein (20–30 µg per lane) were resolved by SDS–PAGE on 10–12% polyacrylamide gels and transferred onto PVDF membranes (0.45 µm pore size).

Membranes were blocked for 1 h at room temperature in 5% nonfat dry milk prepared in TBS containing 0.1% Tween-20 (TBST) and then incubated overnight at 4 °C with primary antibodies specific for phosphorylated NF-κB p65, phosphorylated p38 MAPK, phosphorylated STAT proteins, or corresponding total proteins, diluted in 5% BSA/TBST according to the manufacturers’ recommendations. After washing, membranes were incubated with HRP-conjugated secondary antibodies for 1 h at room temperature.

### Chemokine array

Chemokine profiling was performed using conditioned media collected from human B cells or Bregs. Human B cells were cultured under the indicated conditions, including unstimulated B cells and CpG-stimulated B cells, after which supernatants were harvested and clarified by centrifugation. Where indicated, conditioned media were concentrated prior to analysis.

Chemokine expression was assessed using the Proteome Profiler Human Chemokine Array Kit (R&D Systems, ARY017) according to the manufacturer’s instructions. Array membranes were incubated with conditioned media samples, followed by detection with the supplied biotinylated antibody cocktail and streptavidin–HRP.

Signals were visualized by chemiluminescence and captured using a digital imaging system. Relative chemokine expression was evaluated by comparing signal intensities between experimental conditions. Representative arrays are shown from at least three independent experiments using conditioned media derived from independent human donors.

Protein bands were visualized using enhanced chemiluminescence (ECL). Where indicated, membranes were stripped and reprobed for total protein or housekeeping controls to confirm equal loading. Representative blots shown reflect results obtained from at least three independent experiments using B cells from independent human donors.

### Cytokine Assays

Human IL-10 and TNF-α secreted in the supernatant was measured by ELISA kit (eBioscience, San Diego, CA) by using cytokine-specific pair antibodies according to the manufacturer’s protocol.

### Flow Cytometry

Single-cell suspensions from PBMCs or B cells or T cells were prepared and incubated with the appropriate antibodies at 4 °C for 30 minutes. Then cells were washed and analyzed by a Attune NxT flow cytometer, and data were analyzed by FCS express software. For B cell staining, CD19 (BD Pharmingen™ APC Mouse Anti-Human CD19, cat. #555415), CD25 (BD Pharmingen™ APC Mouse Anti-Human CD25, cat. #567316), CD71 (BD Pharmingen™ PE Mouse Anti-Human CD71, cat. # 555537), CD73 (BD Pharmingen™ BV480 Mouse Anti-Human CD73, cat. #746568). For T cells staining CD3 (BD Pharmingen™ APC Mouse Anti-Human CD3, cat. # 555342), Ki67 (BD Pharmingen™ PE Mouse Anti-Ki-67 Set, cat. # 556027), Fix and Perm buffer (BD Pharmingen™ Fixation/Permeabilization Kit, cat. # 554714), for MDM identification CD14 (BioLegend, APC anti-human CD14 Antibody, cat. # 325608) and CD16 (BioLegend, FITC anti-human CD16, cat. # 980112). Live and dead staining was Tombo Ghost Dye™ Red 780.

### Human RNA-seq sample preparation

Human B cells were isolated from peripheral blood mononuclear cells (PBMCs) obtained from healthy adult donors (n = 3; 2 females, 1 male) using a BioLegend human B cell isolation kit, according to the manufacturer’s instructions. Purity of isolated B cells was routinely >95% as assessed by flow cytometry (CD19⁺).

Purified human B cells were either cryopreserved and used as controls or cultured in complete R10 medium and stimulated under the following conditions: (i) CpG alone (0.1 µM; B-CpG), (ii) purified CD40scFv–IL-21 (100 ng/mL), or (iii) purified CD40scFv–IL-21 (100 ng/mL) in combination with CpG (0.1 µM) to generate IL-10⁺GzmB⁺ regulatory B cells (Bregs). Cells were cultured for 3–6 days, after which cell pellets were harvested for RNA extraction.

### RNA extraction and quality control

Total RNA was extracted from six human B-cell pellets using the Promega Maxwell® RSC simplyRNA Tissue Kit, following the manufacturer’s protocol. RNA concentration was quantified using a Qubit Fluorometer with Broad Range (BR) reagents. RNA integrity was assessed using an Agilent TapeStation with High Sensitivity RNA ScreenTape, and only samples with RNA integrity number (RIN) ≥7 were used for library preparation.

### Library preparation and sequencing

Poly(A)-selected, strand-specific RNA-seq libraries were prepared using the Illumina TruSeq Stranded mRNA Library Prep Kit. Library size distribution and quality were confirmed using the Agilent TapeStation with D1000 ScreenTape. Sequencing was performed on an Illumina NovaSeq X platform using paired-end 150 bp reads (PE150), generating approximately 20 million paired-end reads per sample.

### Computational processing and differential expression analysis

Raw FASTQ files were subjected to quality control using FastQC. Adapter sequences and low-quality bases were removed using Trim Galore. Cleaned reads were aligned to the human reference genome (GRCh38) using STAR. Gene-level read counts were generated using HTSeq.

Differential gene expression analysis was performed using DESeq2 with a design formula of ∼ condition (Bregs vs B-CpG vs unstimulated B cells vsB-CD40scFv-IL21). Statistical significance was defined as an adjusted p-value (padj) < 0.05 using Benjamini–Hochberg false discovery rate correction. For visualization, variance-stabilizing transformed (VST) counts were used to generate principal component analysis (PCA) plots and heatmaps.

### Exploratory and functional enrichment analyses

Exploratory transcriptomic analyses and gene-set visualizations were performed using iDEP (v0.96). Normalized counts were log1p-transformed and z-scored for heatmap visualization. Functional enrichment and Gene Ontology analyses were conducted using DAVID (v2024q4) and Graphite Web to identify immune-regulatory pathways associated with the Breg transcriptional program.

## Data availability

Raw sequencing data (FASTQ files), processed gene-level count matrices, and associated metadata will be deposited in the NCBI Gene Expression Omnibus (GEO) upon manuscript acceptance.

### Human T-cell suppression assay

For human T-cell suppression assays, PHA-activated T cells were co-cultured with autologous B-cell populations under the indicated conditions. Regulatory B cells (Bregs) were generated by stimulating purified human B cells with CD40scFv–IL-21 (100 ng/mL) in combination with CpG ODN2006 (0.1 µM) for 3-6 days. Control conditions included unstimulated B cells and CpG-stimulated B cells (B-CpG). Following stimulation, B cells were washed and added to activated T cells at the indicated ratios.

After 3 days of co-culture, T-cell proliferation was assessed by intracellular Ki-67 staining within live CD3⁺ T cells using flow cytometry. Data were acquired on an Attune NxT flow cytometer and analyzed using FCS Express software. Suppressive activity was calculated relative to PHA-stimulated T cells cultured in the absence of B cells. Experiments were performed using PBMCs from five independent human donors.

### *In vivo* xenogeneic graft-versus-host disease (GVHD) model

To evaluate the in vivo immunomodulatory activity of Bregs generated using CD40scFv–IL-21, a xenogeneic graft-versus-host disease (GVHD) model was employed. NOD-SCID-IL2Rγnull (NSG) mice were sublethally irradiated with 2 Gy total body irradiation. Twenty-four hours later, mice received 6 × 10⁶ human T cells administered intravenously, either alone (PBS control) or premixed with autologous Bregs at the indicated ratio prior to infusion. Mice were monitored longitudinally for survival as the primary readout of disease progression. Animals were observed daily and euthanized upon reaching humane endpoints in accordance with institutional animal care guidelines. Survival curves were generated to assess the impact of Bregs on T cell–mediated disease severity.

## Statistical analysis

Quantitative data are presented as mean ± standard error of the mean (SEM) unless otherwise indicated. All in vitro experiments were performed using biological replicates derived from independent human donors, as specified in the figure legends.

For flow cytometry–based phenotypic analyses and cytokine quantification (Figures 2, 5, and 6), data represent at least five independent biological replicates. Western blot and signaling analyses (Figures 1 and 3) are representative of a minimum of three to four independent experiments performed using cells from independent donors. Normality was not assumed a priori due to limited sample sizes. For comparisons involving two experimental groups, statistical significance was assessed using an unpaired, two-tailed Student’s t-test when data met assumptions of normality. For comparisons involving three or more groups, one-way analysis of variance (ANOVA) followed by Tukey’s multiple-comparison post hoc test was applied (Figures 2B, 2C, and 6). For non-parametric data, including T-cell suppression assays (Figure 5), statistical significance was evaluated using the Mann–Whitney U test.

**Figure 1.**
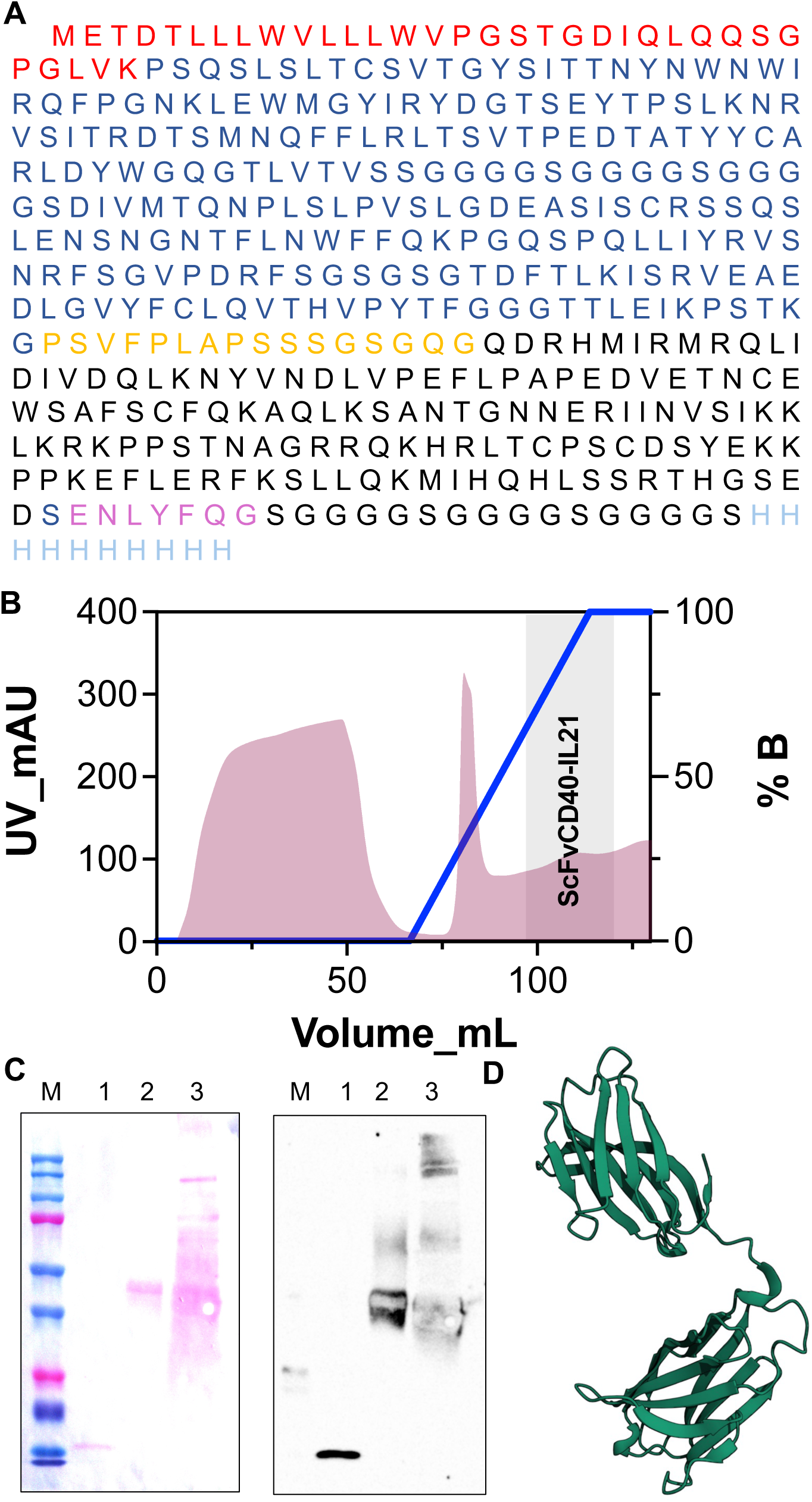
Design, purification, and structural modeling of the CD40scFv–IL-21 fusion protein. **(A)** Amino acid sequence of the CD40scFv–IL-21 fusion protein (441 amino acids). The signal peptide is shown in red, the CD40-specific scFv in blue, the linker region in orange, and the IL-21 domain in black. A C-terminal spacer containing a TEV protease recognition site (ENLYFQG), followed by an additional spacer and an 8×His affinity tag, is included to facilitate purification and optional tag removal. **(B)** Representative FPLC chromatogram of Ni–NTA affinity purification. Conditioned medium from stable transduced HEK293T cells was pH-adjusted and loaded onto a 5 mL Ni–NTA column equilibrated in binding buffer. Elution was performed over 10 column bed volumes using a gradient to 100% buffer B. The dark shaded grey region indicates the pooled fractions containing purified CD40scFv–IL-21. **(C)** Ponceau S staining and Western blot analysis of purified CD40scFv–IL-21. The fusion protein was detected using an anti-human IL-21 antibody. Lane M, molecular weight markers; lane 1, recombinant human IL-21 (200 ng); lane 2, purified CD40scFv–IL-21; lane 3, conditioned media from transduced 293T cells. The expected molecular weight is 45.4 kDa. **(D)** Predicted three-dimensional structure of CD40scFv–IL-21 generated from the amino acid sequence using the Phyre2 protein modeling server [46].

**Figure 2.**
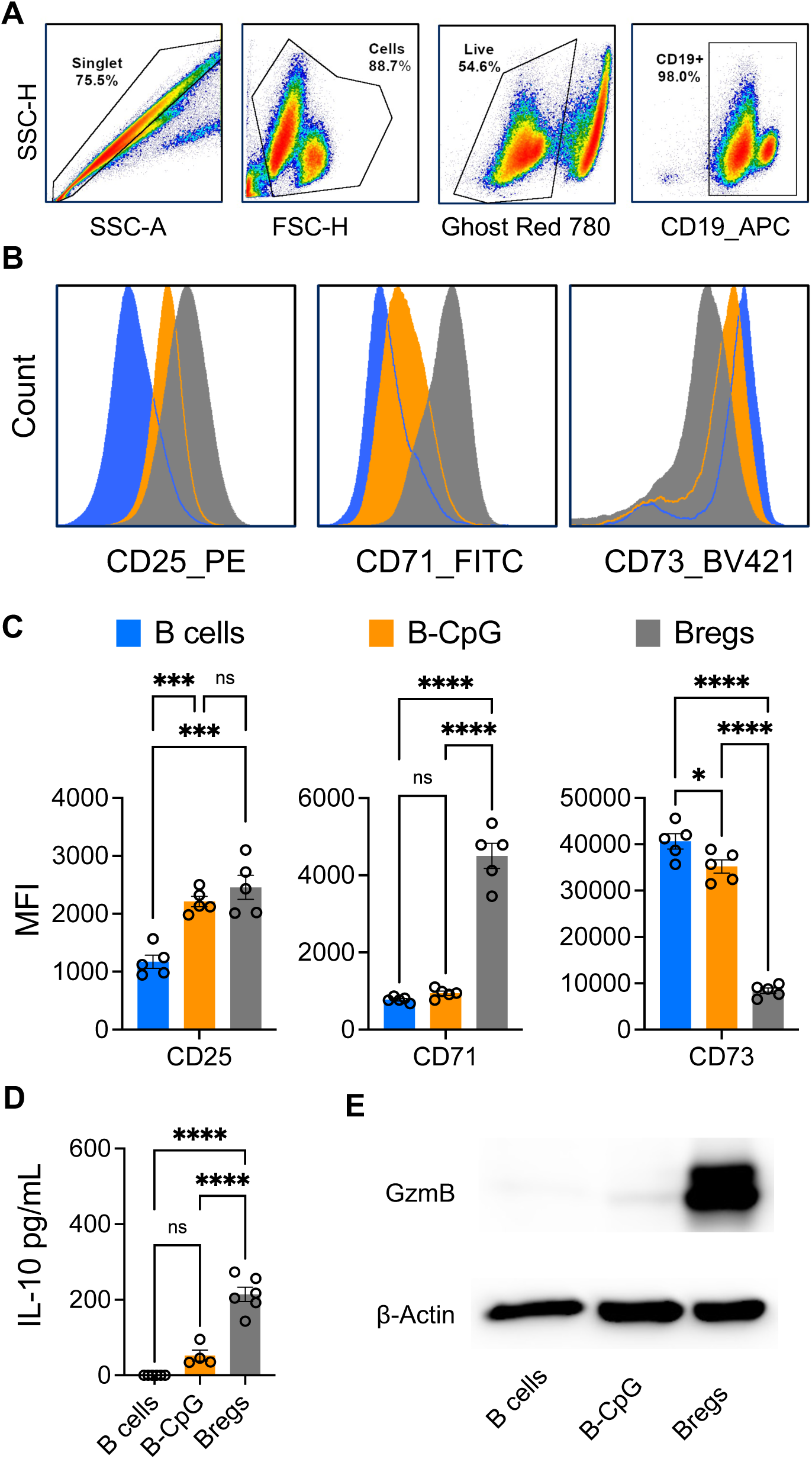
Phenotypic and functional characterization of B cells, B-CpG, and Bregs cells. (**A**) Flow cytometry analysis showing the gating strategy and expression of CD25, CD71, and CD73 in B cells (blue), B-CpG (orange), and Bregs cells (gray). (**B**) Bar graphs showing median fluorescence intensity (MFI) determined by flow cytometry in (A). Bars represent the mean ± SEM of CD25, CD71, and CD73 expression from five independent samples. (**C**) IL-10 expression in conditioned media from B cells, B-CpG, and Bregs cells. Bar graphs represent mean ± SEM values from 5 different samples. (**D**) Western blot analysis of Granzyme B (GzmB) expression in B cells, B-CpG, and Bregs. β-actin was used as a housekeeping protein and loading control. Statistical analysis for panels (**B**) and (**C**) was performed independently for each marker using one-way ANOVA followed by Tukey’s multiple-comparison test. Statistical significance is indicated by asterisks as follows: * p < 0.05; ** p < 0.01; *** p < 0.001; **** p < 0.0001.

**Figure 3.**
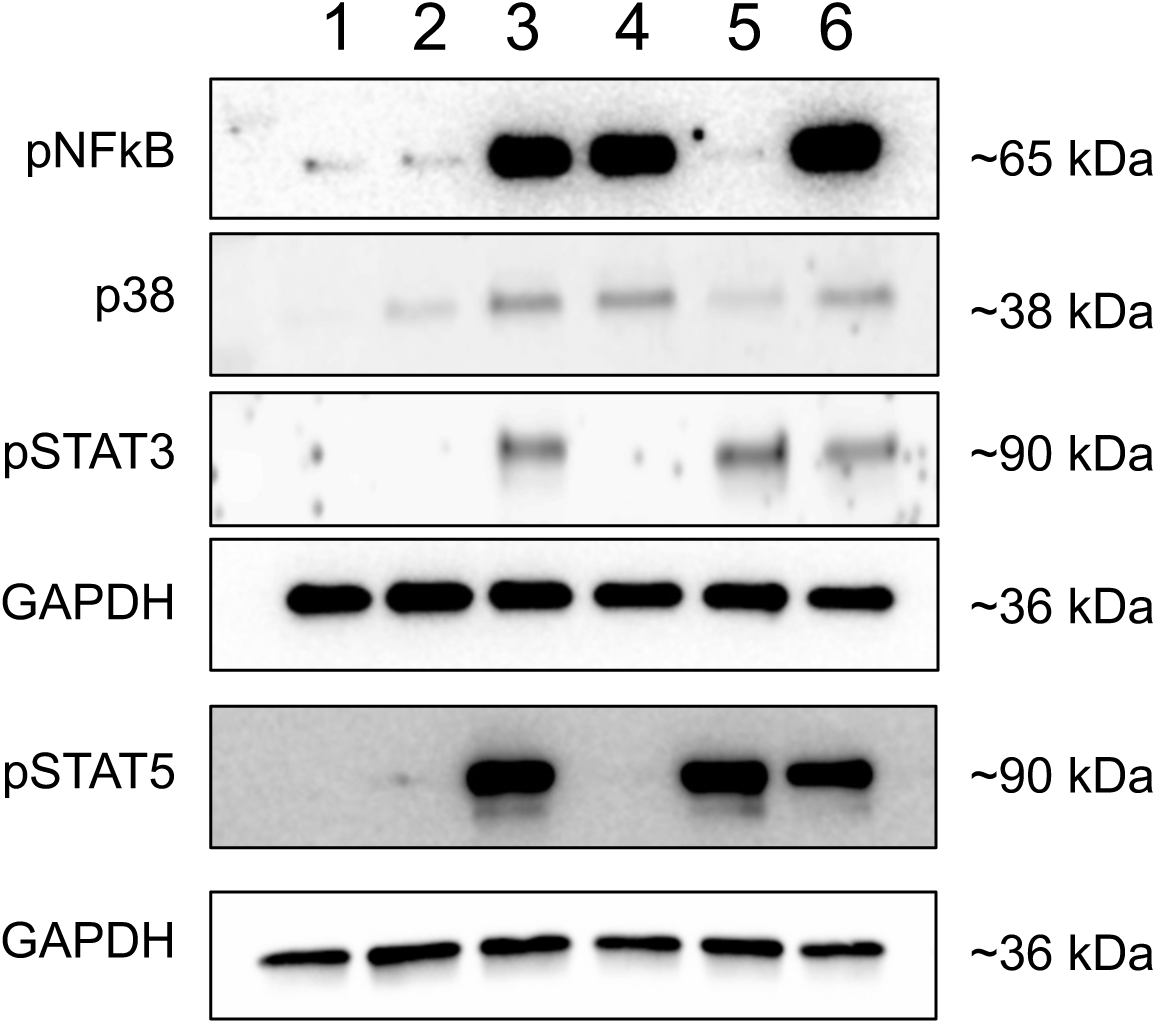
CD40scFv–IL-21 induces NF-κB activation consistent with agonistic CD40 signaling and preserves IL-21–dependent signaling pathways. Human peripheral blood B cells were stimulated as indicated with CD40scFv–IL-21, CpG, or control conditions. Lane assignments were as follows: 1, unstimulated control; 2, CpG; 3, CpG, CD40 ligand (CD40L) and IL-21; 4, CpG and CD40scFv–IL-21; 5, CpG and IL-21. Whole-cell lysates were analyzed by immunoblotting for phosphorylated NF-κB p65 (p-NF-κB) and downstream signaling intermediates. IL-21–dependent signaling was preserved under these conditions. GAPDH and β-actin was used as a loading control. Representative blots from independent experiments are shown. Blots shown are representative of more than four independent experiments performed using B cells isolated from independent human blood donors.

**Figure 4.**
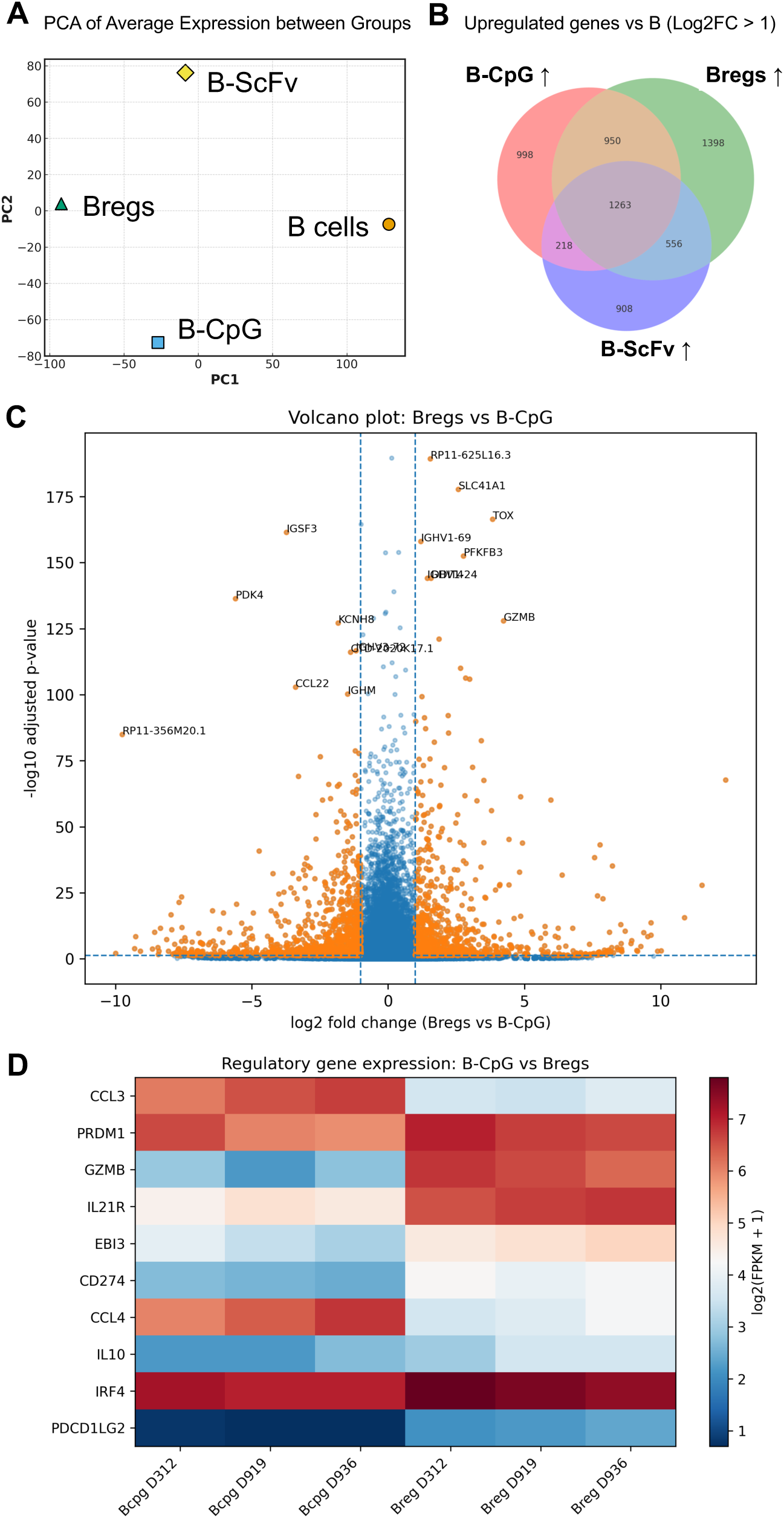
Transcriptomic profiling of human peripheral blood B cells identifies a regulatory gene program induced by CD40scFv–IL-21 and CpG. **(A)** Principal component analysis (PCA) of RNA-seq data from purified human peripheral blood B cells cultured under four conditions: unstimulated B cells, CpG-stimulated B cells (B-CpG), CD40scFv–IL-21–stimulated B cells (B-ScFv), and B cells stimulated with the combination of CD40scFv–IL-21 and CpG (Bregs). PCA demonstrates clear separation among conditions, indicating distinct global transcriptional states induced by single versus combined stimulation. **(B)** Venn diagram showing the number of genes upregulated (log₂ fold change > 1) relative to unstimulated B cells in B-CpG, B-ScFv, and Bregs. The overlap highlights both shared and condition-specific transcriptional responses, with Bregs exhibiting a distinct set of upregulated genes not observed with single-stimulus activation. **(C)** Volcano plot depicting differential gene expression between Bregs and B-CpG cells. The x-axis represents log₂ fold change (Bregs vs B-CpG), and the y-axis represents −log₁₀ adjusted p-value. Genes significantly upregulated in Bregs include regulatory and effector-associated transcripts such as *IL10, GZMB, CD274 (PD-L1), PDCD1LG2 (PD-L2)*, and *EBI3*, whereas CpG-associated inflammatory and chemokine-related genes are less prominent under combined stimulation. **(D)** Heat map showing expression of selected regulatory, checkpoint, transcriptional, and chemokine-associated genes comparing B-CpG and Bregs across three independent donors (D312, D919, D936). Expression values are displayed as log₂(FPKM + 1) and visualized using a red–blue diverging color scale, with red indicating higher relative expression and blue indicating lower expression. The heat map reveals coordinated upregulation of regulatory and effector-associated genes (*IL10, GZMB, CD274, PDCD1LG2, EBI3, PRDM1, IRF4*) in Bregs, whereas B-CpG cells show relatively higher expression of chemokines such as *CCL3* and *CCL4*.

**Figure 5.**
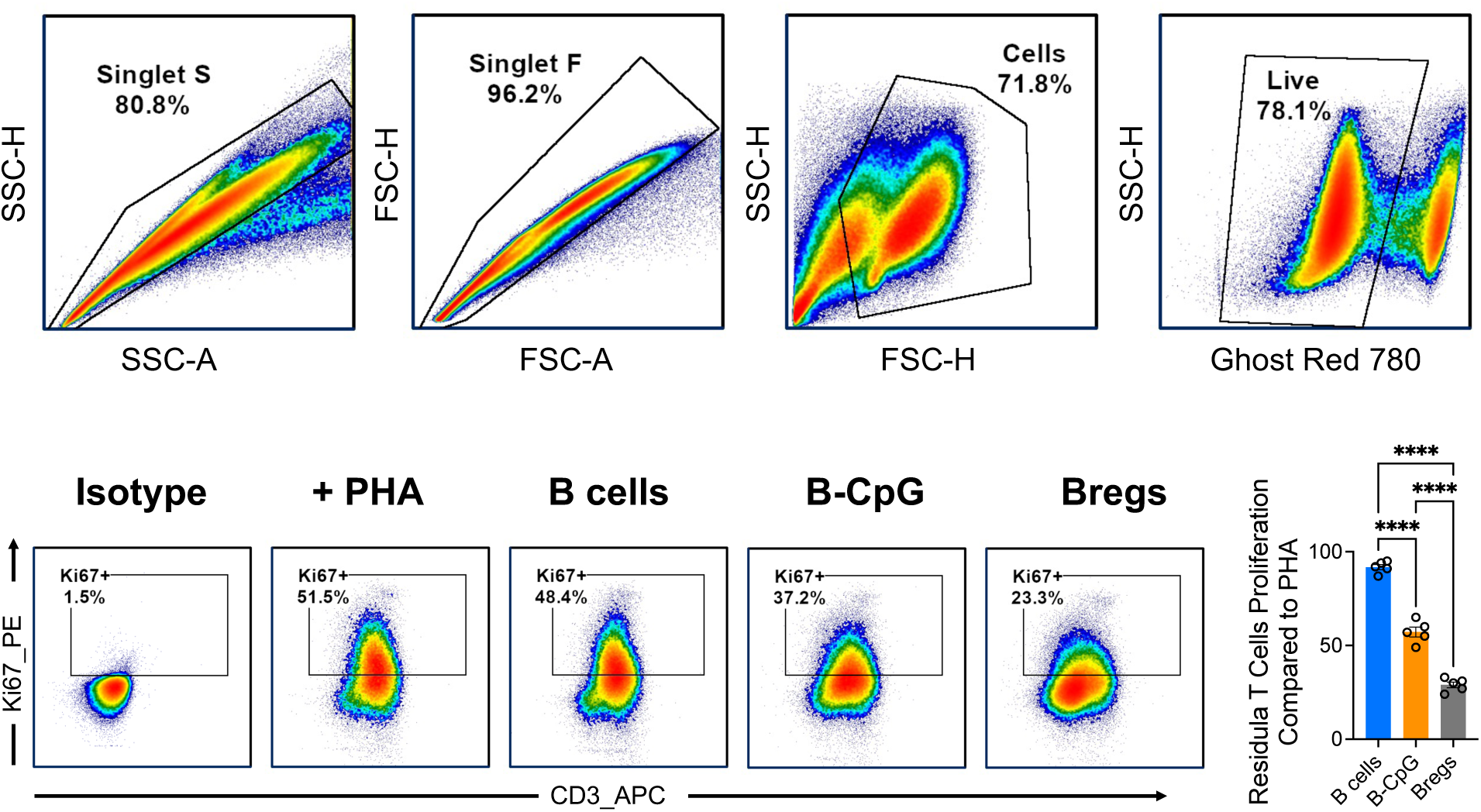
**T cells suppression assay**. (**A**) Flow cytometry gating strategy. Singlets were identified based on sized and forward scatter, followed by size and granularity (FSC/SSC) gating and exclusion of dead cells using a live/dead dye. (**B**) Ki-67 staining within gated CD3⁺ T cells under the indicated conditions: isotype control, +PHA, co-culture with B cells, B-CpG cells, or Bregs. (**C**) Bar graphs of T cells proliferation suppression, statistical analysis has been carried out with 5 biological replicates. Statistical analysis was performed using the Mann–Whitney U test with 5 biological replicates. Data are presented as mean ± SEM. Statistical significance is * p < 0.05; ** p < 0.01; *** p < 0.001; **** p < 0.0001.

RNA-sequencing differential expression analysis (Figure 4) was performed using DESeq2, with p-values adjusted for multiple testing using the Benjamini–Hochberg false discovery rate (FDR) method. Genes were considered significantly differentially expressed when the adjusted p-value was < 0.05 and the absolute log₂ fold change exceeded 1, unless otherwise stated.

All statistical analyses were performed using GraphPad Prism software. Statistical significance was defined as p < 0.05. Levels of significance are indicated in the figures as follows: p < 0.05 (), p < 0.01 (), p < 0.001 (), and p < 0.0001 (****).

## Results

### Design, expression and purification of CD40ScFv-IL21

The CD40scFv–IL-21 fusokine was created by fusing the single chain variable portion of a CD40 antibody DNA sequence, preceded by its signal peptide, followed by the sequence encoding for a short linker and the mature human IL-21 without signal peptide. A C-terminal spacer containing a TEV protease recognition site, followed by an additional spacer and an 8×His affinity tag, was included to facilitate purification and optional tag removal. The final sequence has a length of 1,273 nucleotides and encodes for a single polypeptide of 423 amino acids (Fig. 1A).

To generate the CD40scFv–IL-21 fusion protein human 293T cell line stably expressing CD40scFv–IL-21 was cultured in suspension using FreeStyle™ 293 Expression Medium. Cells were expanded in shake flasks under serum-free conditions and maintained at high viability (>90%) throughout the culture period. Conditioned media were collected and supernatants containing CD40scFv–IL-21 were subsequently used for downstream purification.

The conditioned medium was loaded onto a 5 mL Ni–NTA affinity column (GE Healthcare) equilibrated in binding buffer consisting of 20 mM sodium phosphate, 300 mM NaCl, and 5 mM imidazole, pH 7.4. After washing to remove unbound and weakly bound proteins, CD40scFv–IL-21 was eluted using a linear gradient from 0 to 100% buffer B (20 mM sodium phosphate, 300 mM NaCl, 500 mM imidazole, pH 7.4) over 10 column bed volumes. Fractions corresponding to the major elution peak, indicated by the shaded gray region of the chromatogram, were pooled, concentrated to a final volume of 2 mL, and subsequently loaded onto a PD-10 desalting column equilibrated in PBS for buffer exchange prior to downstream analyses (Fig. 1B).

The molecular weight of the purified protein, predicted to be approximately 45 kDa, was confirmed by Western blot analysis using an anti-human IL-21 antibody. Ponceau S staining of the same samples did not reveal additional protein bands, indicating a high degree of purity of the CD40scFv–IL-21 preparation (Fig. 1C). Computer analysis and structure prediction based on amino acid sequence reveal the immunoglobin-like domains of CD40ScFv and IL21 (Fig. 1D). CD40scFv–IL-21 quantification was carried out by human IL-21 ELISA.

### Generation of IL-10⁺GzmB⁺ B cells (Bregs)

Human B cells isolated from peripheral blood mononuclear cells were stimulated with purified CD40scFv–IL-21 100 ng/mL in combination with 0.1 μM CpG to generate IL-10⁺GzmB⁺ B cells (Bregs), or with 0.1 μM CpG alone (B-CpG), with unstimulated B cells serving as controls. After 3–6 days of culture, cells were analyzed by flow cytometry to assess B-cell purity (CD19) and the expression of CD25, CD71, and CD73 (Fig. 2A).

Flow cytometric analysis revealed that stimulation with CD40scFv–IL-21 and CpG induced a distinct B-cell phenotype characterized by increased expression of the activation markers CD25 and CD71 and reduced expression of CD73 compared with both unstimulated B cells and B-CpG–treated cells (Fig. 2A,B). Quantitative analysis of median fluorescence intensity (MFI) showed significant differences among the three groups for CD25, CD71, and CD73 expression (one-way ANOVA; CD25: *P* = 0.0001; CD71: *P* < 0.0001; CD73: *P* < 0.0001), with Tukey’s multiple-comparison test confirming that Bregs cells differed significantly from both B cells and B-CpG cells for all three markers (Fig. 2B,C).

Given the central role of IL-10 in regulatory B-cell function, IL-10 secretion was assessed in conditioned media by ELISA. While CpG stimulation alone induced a modest increase in IL-10 production that did not reach statistical significance compared with unstimulated B cells, stimulation with CD40scFv–IL-21 in combination with CpG resulted in a marked and statistically significant increase in IL-10 secretion (*P* < 0.0001; one-way ANOVA with Tukey’s post hoc test) (Fig. 2D). IL-10 was not detected in supernatants from unstimulated B cells (Fig. 2D).

Expression of Granzyme B (GzmB) was assessed by Western blot analysis (Fig. 2E). GzmB protein was readily detected in B cells stimulated with the combination of CD40scFv–IL-21 and CpG, whereas little to no GzmB expression was observed in unstimulated B cells or in cells treated with CpG alone (Fig. 2E). β-actin was used as a loading control.

### CD40scFv–IL-21 activates canonical CD40 signaling and NF-κB phosphorylation in human B cells

To determine whether the CD40-targeting scFv component of the CD40scFv–IL-21 fusion protein functions as a bona fide agonist, we assessed activation of downstream CD40 signaling pathways in stimulated B cells. Engagement with CD40scFv–IL-21 resulted in robust phosphorylation of NF-κB p65, a defining feature of canonical CD40 signaling, at levels comparable to those induced by CD40 ligand stimulation (Fig. 3). In addition to NF-κB activation, CD40scFv–IL-21 stimulation induced p38 MAPK phosphorylation, further supporting effective engagement of TRAF-dependent CD40 signaling cascades. These signaling responses demonstrate that the anti-CD40 scFv functions as an agonistic ligand, rather than merely a targeting or blocking moiety.

In parallel, IL-21–dependent STAT phosphorylation was readily detected following stimulation with CD40scFv–IL-21, indicating that fusion of IL-21 to the CD40-binding scFv does not compromise IL-21 receptor engagement or downstream JAK–STAT signaling. Notably, STAT activation was not observed following CD40 engagement alone, consistent with pathway specificity and confirming that STAT signaling is driven by the IL-21 component of the fusion protein.

Collectively, these data demonstrate that CD40scFv–IL-21 simultaneously activates CD40-associated NF-κB and p38 pathways while preserving IL-21–mediated STAT signaling (Fig. 3).

### Transcriptomic analysis of human peripheral blood B cells identifies a regulatory program induced by CD40scFv–IL-21 and CpG

To define the transcriptional identity of human regulatory B cells induced by CD40scFv–IL-21 and CpG, RNA sequencing was performed on purified peripheral blood B cells cultured under four conditions: unstimulated B cells, CpG-stimulated B cells (B-CpG), CD40scFv–IL-21–stimulated B cells, and B cells stimulated with the combination of CD40scFv–IL-21 and CpG (Bregs), using samples from three healthy donors (two female and one male). Principal component analysis of average gene expression revealed clear separation among all four conditions, demonstrating that combined CD40scFv–IL-21/CpG stimulation induces a transcriptional state distinct from resting B cells and from either stimulus alone (Fig. 4A). Notably, Bregs segregated from B-CpG cells along both principal components, indicating that the combined stimulus does not merely enhance CpG-driven activation but instead promotes a distinct transcriptional program.

To further characterize transcriptional differences, differentially expressed genes (DEGs) were identified relative to resting B cells and visualized using a Venn diagram (Fig. 4B). This analysis revealed both shared and condition-specific gene expression changes across B-CpG, CD40scFv–IL-21, and Breg conditions, with a substantial subset of genes uniquely upregulated in Bregs. These data indicate that CD40scFv–IL-21/CpG stimulation induces a gene expression program that is only partially overlapping with CpG-induced activation and includes a distinct set of transcripts not observed under single-stimulus conditions.

Direct comparison of Bregs and B-CpG cells identified extensive transcriptional differences, which are visualized in a volcano plot (Fig. 4C). Numerous genes were significantly upregulated in Bregs relative to B-CpG cells, reflecting broad transcriptional divergence between these two activated states. Among the genes showing increased expression in Bregs were transcripts associated with regulatory or effector-associated functions, including *IL10, GZMB, CD274* (PD-L1), *PDCD1LG2* (PD-L2), *EBI3, PRDM1*, and *IRF4*. In contrast, genes preferentially expressed in B-CpG cells included chemokines such as *CCL3* and *CCL4*, consistent with a CpG-driven inflammatory activation profile. Comprehensive lists of the top 100 up-and down-regulated genes identified in comparisons of Bregs with resting B cells and with B-CpG cells are provided in Supplementary Tables S1 and S2, respectively.

To assess donor-level consistency and coordinated expression patterns, a focused heat map was generated comparing B-CpG and Bregs across individual donors (Fig. 4D). Expression values are displayed as log₂(FPKM + 1) and visualized using a red–blue color scale, with red indicating higher relative expression and blue indicating lower expression. This analysis demonstrated consistent upregulation of regulatory and effector-associated genes, including *IL10, GZMB, CD274, PDCD1LG2, EBI3, PRDM1*, and *IRF4*, across donors in the Breg condition, whereas B-CpG cells showed relatively higher expression of inflammatory chemokines such as *CCL3* and *CCL4*. Together, these data support the conclusion that combined CD40scFv–IL-21 and CpG stimulation induces a reproducible regulatory transcriptional program that is distinct from CpG stimulation alone.

### T-cells suppression assay

To evaluate the suppressive capacity of regulatory B cells (Bregs), in vitro T-cell suppression assays were performed using human PBMCs stimulated with PHA. Proliferation of responder T cells was assessed by Ki-67 expression within gated CD3⁺ T cells following 3 days of co-culture. Flow cytometric analysis was conducted using a sequential gating strategy consisting of singlet discrimination, size and granularity (FSC/SSC) gating, and exclusion of non-viable cells using a live/dead dye (Fig. 5A).

Under PHA stimulation, T cells displayed robust proliferative responses as expected (Fig. 5B). Co-culture with unstimulated B cells did not significantly alter T-cell proliferation, indicating minimal suppressive activity under these conditions. In contrast, CpG-stimulated B cells (B-CpG) induced a modest but significant reduction in T-cell proliferation relative to B cells alone (Fig. 5C).

Notably, co-culture with Bregs resulted in a marked suppression of T-cell proliferation, significantly exceeding the inhibitory effects observed with either B cells or B-CpG cells (Fig. 5C). This suppressive effect was consistently observed across five independent human blood donors and was not associated with reduced T-cell viability, indicating that Breg-mediated suppression reflects functional inhibition rather than cytotoxicity.

### Polarization of peripheral myeloid cells with IL-10^+^GzmB^+^ B cells

Human peripheral blood mononuclear cells (PBMCs) were separated by density centrifugation using Ficoll. CD14 beads were used to positively select monocytes according to the manufacturer’s protocol. The purity of the cells was checked by flow cytometry after each isolation, and the typical purity for monocytes is >98%. Monocyte-derived macrophages (MDM) were differentiated for 5 days as described previously. Condition media of unstimulated B cells, B-CpG or Bregs cells were added to the MDM for 24 h and LPS was added to a concentration of 100 ng/mL for 24 h in fresh media after which MDM condition media was tested for the expression of TNF-α. MDM treated with BREGS cells condition media showed the lowest expression in TNF-α compared to MDM treated with condition media from unstimulated cells (B sup, p=0.0008) or B-CpG condition media (B-CpG sup, p= 0.006) (Fig. 6). Condition media from B-CpG cells did not decrease the expression of TNF-α similarly to what observed from unstimulated B cells condition media (Fig. 6).

**Figure 6.**
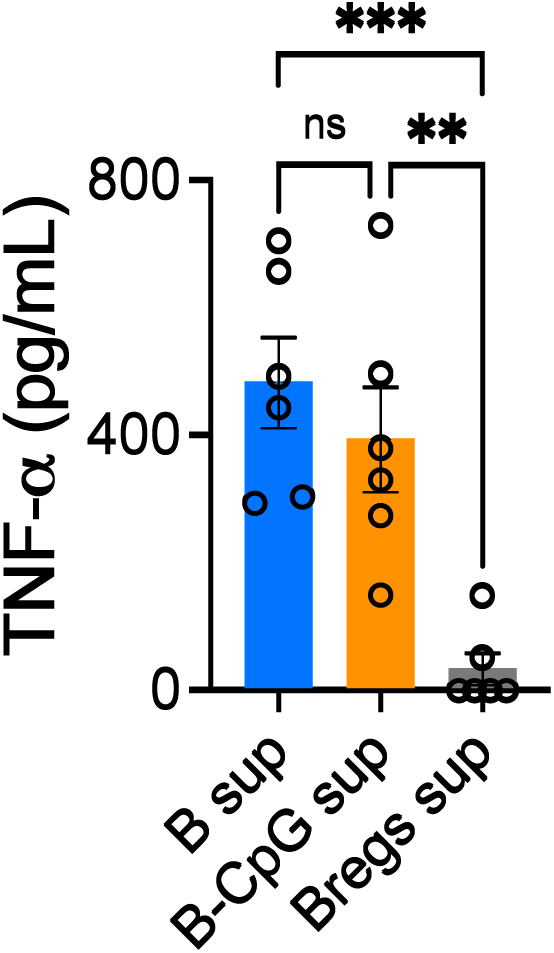
Modulation of monocyte-derived macrophages phenotype. Expression of TNF-α, upon cultured of CD14^+^ monocyte-derived macrophages LPS stimulated with B cells supernatant (B sup), B-CpG supernatant (B-CpG sup) or Bregs supernatant (Bregs sup). Cytokine expression was measured by ELISA. Data are expressed as the mean – SEM and one-way ANOVA followed by Tukey’s multiple-comparison test. Statistical significance is * p < 0.05; ** p < 0.01; *** p < 0.001; **** p < 0.0001.

### Bregs delay T cell–mediated xenoGVHD disease progression in NSG mice

To assess the impact of regulatory B cells (Bregs) on T cell–mediated disease progression in vivo, NSG mice were sublethally irradiated with 2 Gy and subsequently administered 6 × 10⁶ human T cells, either premixed with Bregs or co-administered with PBS as control. Following adoptive transfer, mice were monitored longitudinally, and survival was used as the primary readout to capture overall disease progression (Fig. 7). Mice receiving T cells alone exhibited a progressive decline in survival over time, consistent with the development of T cell–driven pathology in this model. In contrast, mice administered T cells premixed with Bregs demonstrated a delayed onset of mortality and improved overall survival relative to PBS controls. Survival curves revealed a clear temporal separation between groups, indicating that the presence of Bregs modulated the course of disease progression following T-cell transfer.

**Figure 7.**
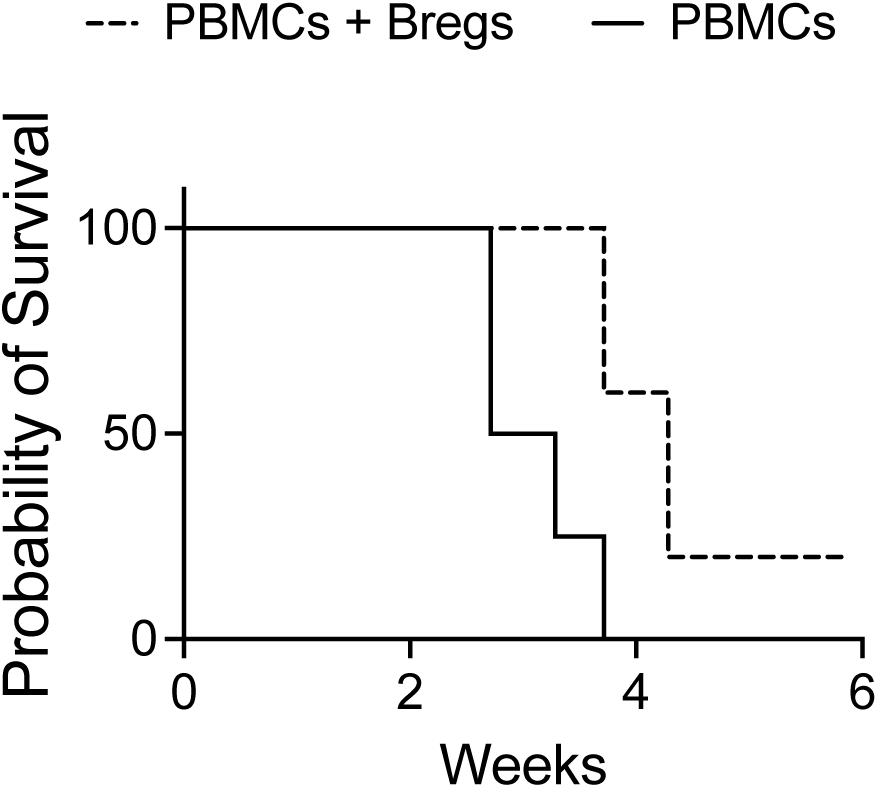
Regulatory B cells delay T cell–mediated disease progression in irradiated NSG mice. NOD-SCID-IL2Rγnull (NSG) mice were sublethally irradiated and intravenously injected with 6 × 10⁶ human T cells either alone (T cells only) or premixed with autologous IL-10⁺GzmB⁺ regulatory B cells (Bregs) generated ex vivo using CD40ScFv–IL-21 and CpG stimulation. Mice were monitored longitudinally, and overall survival was used as the primary endpoint to assess T cell–mediated disease progression. Mice receiving T cells alone exhibited progressive mortality consistent with xenogeneic T cell–driven pathology, whereas co-administration of Bregs significantly delayed disease onset and improved survival. Survival curves were generated using the Kaplan–Meier method, and statistical significance was determined using the log-rank (Mantel–Cox) test. Data are representative of at least three independent experiments with n = 5 mice per group.

No additional clinical scoring was applied, and survival was intentionally used as an integrative endpoint to reflect cumulative disease burden rather than discrete pathological features. Together, these data suggest that Bregs attenuate T cell–mediated disease severity in irradiated NSG mice, as reflected by prolonged survival (Fig. 7).

## Discussion

Regulatory B cells (Bregs) are critical mediators of immunological tolerance, exerting immunosuppressive effects through the secretion of polarizing cytokines and by limiting the expansion of pathogenic T cells and other pro-inflammatory lymphoid and myeloid populations[28, 29]. In murine systems, multiple protocols have been established to convert B cells into IL-10–competent regulatory populations [30], and adoptive transfer of such manufactured Bregs has shown therapeutic benefit in several inflammatory disease models [31–34]. In human, the frequency of Bregs in blood is less than 1-2% [35] and both numerical and functional defects in this compartment have been reported in autoimmune diseases [36]. The vexing obstacle in translational use of Bregs for humans, as an investigational pharmaceutical, is that conversion of human B-cells to Bregs is unresponsive to techniques developed for the genesis of mouse Bregs [25, 26, 37, 38]. There are no practical methods for the *ex vivo* GMP expansion of human regulatory B cells that enable first in human clinical study which likely explains the absence of investigative clinical trials with this cell therapy platform.

In the present study, we describe a novel composition of matter that enables rapid and robust conversion of human peripheral B cells into B cells with a regulatory phenotype *in vitro*. This bifunctional fusion protein, CD40scFv–IL-21, combines an agonistic anti-human CD40 single-chain variable fragment with human IL-21. When used in combination with CpG, this synthetic leukine converts human B cells into regulatory B cells (Bregs) competent for IL-10 and granzyme B (GzmB) expression within less than 72 hours. Importantly, a critical translational requirement was met by demonstrating that CD40scFv–IL-21 can be successfully purified from producer cell–conditioned media to homogeneity. Affinity-based purification yielded a biochemically intact protein free of detectable contaminating conditioned-media proteins. Functional assays confirmed that the purified fusion protein retained full CD40 agonistic activity and IL-21–dependent signaling, establishing that the observed biological effects are attributable to the defined molecular entity rather than to accessory factors. This biochemical robustness and manufacturability strongly support the feasibility of GMP-scale production.

The enhanced stability of the bifunctional CD40scFv–IL-21 protein relative to recombinant IL-21 or the CD40-binding domain alone is a key translational advantage. Importantly, the CD40ScFv functions as a true agonist, engaging CD40 in a manner that mimics physiological CD40L signaling and triggers downstream activation pathways. Prior approaches to induce human Bregs relied on co-culture with CD40L-expressing feeder cells [39, 40], making the use of these technologies less translational for GMP manufacturing at scale.

IL-21 is the newest member of the common γ-chain family of cytokines and has been shown to have a pleiotropic function on many cells, including B cells [41]. While IL-21 alone can synergize with TLR signaling to induce growth arrest or apoptosis [42], we observed that, when delivered in conjunction with CD40 agonism, IL-21 instead promoted robust B-cell expansion. This observation is consistent with prior reports showing that CD40 signaling converts IL-21 into a co-stimulatory signal that enhances cell cycling and proliferation [42]. Moreover, IL-21 induces the expression of granzyme B (GzmB), a main member of the serine proteinase superfamily, as detected by transcriptome analysis. GzmB-RNA from Bregs cells is one of the top 25 upregulated differentially expressed gene compared to the one observed in B-CpG cells. Western blotting analysis, using anti human GzmB antibodies, confirmed the present at a protein level of GzmB in the extract of Bregs cells. It is widely accepted that B cell derived GzmB exerts a non-cytotoxic immunoregulatory effect of target cell surface component (e.g., TCR) [43, 44]. Inhibition of T cells proliferation by Bregs involved transfer of GzmB and degradation of the ζ-chain component of the T cell receptor without induction of cell apoptosis [44]. Bregs cells are competent to inhibit the proliferation of PHA-activated T cells, both in autologous and allogeneic settings, without inducing significant cell dead as determined by flow cytometry analysis.

Despite accumulating data describing the role of cytokine and protein secreted by Bregs subset, there is a lot of heterogeneity about the phenotypic surface fingerprinting of human Bregs. By using flow cytometry analysis, we could identify 3 surface markers that were significantly differently expressed on Bregs cells compared to B cells. Bregs cells showed increased expression of CD25, CD71 and lower expression of CD73 compared to B cells. Interesting, CpG alone marginally downregulated the expression of CD73 as reported in literature [45], whereas the combination of CD40scFv–IL-21 and CpG downregulate the expression of CD73 even further.

Identification of the underlying gene expression changes between Bregs cells and resting B cells or B-CpG cells provided valuable insight into the biological changes leading to the regulatory phenotype in B cells. The study described here showed 283 genes to be differentially expressed between Bregs cells, B cells and B-CpG cells. Within the 283 genes that were differentially expressed, there were several strong candidates’ genes for further study and validation, specifically the one associated with immune regulation. Indeed, gene expressing profiling reveals an upregulation in immune associated genes in Bregs cells compared to B-CpG cells. Negative regulation of T cell activation, leukocyte activation and inflammatory response were the most differentially expressed genes involved in biological processes of Bregs cells compared to B-CpG cells.

We have also developed an *in vitro* potency assay predicated on the ability of Bregs cells to polarize macrophages to an M2 functionality *in vitro*. Our findings reveal an anti-inflammatory B cell/myeloid cell axis, wherein Bregs cells can induce anti-inflammatory myeloid cell responses, including decreased secretion of TNF-α in MDM.

Beyond *in vitro* potency, we evaluated the functional relevance of Bregs cells *in vivo* using a xenogeneic graft-versus-host disease (GVHD) model. NSG mice were irradiated and administered human T cells either alone or pre-mixed with Bregs, and survival was monitored as an integrated measure of disease progression. Co-administration of Bregs cells significantly prolonged survival compared to T cells alone, demonstrating effective suppression of pathogenic T-cell–mediated disease *in vivo*. These findings indicate that BREGS cells retain immunomodulatory activity within a complex inflammatory environment and provide proof-of-concept for their therapeutic potential in T-cell–driven immune pathology.

In summary, our observations and experiments have validated that CD40ScFv-IL21/CpG will convert blood B cells into Bregs cells with a regulatory phenotype within 72 hours. From a pragmatic cell manufacturing perspective, a single volume leukapheresis from a human subject typically yields 1-2 x 10^9^ B-cells, which following enrichment and subsequent 72-hour CD40ScFv-IL21/CpG conversion in vitro provides a clinically enough Bregs for cryobanking and latter investigative use. The development of Bregs cells with a potent regulatory phenotype, furthers the rationale of personalized Bregs adoptive therapy for immune disorders.

## Acknowledgements and disclosures

This work was supported by a University of Wisconsin in Madison Foundation grant and a provisional patent application for CD40ScFv-IL21/CpG has been filed and assigned to WARF (Wisconsin Alumni Research Foundation).

